# Selective Inhibition of Interleukin-2 Inducible T Cell Kinase (ITK) Enhances Anti-Tumor Immunity in Association with Th1-skewing, Cytotoxic T cell Activation, and Reduced T Cell Exhaustion

**DOI:** 10.1101/2023.07.05.547822

**Authors:** Lih-Yun Hsu, James T Rosenbaum, Erik Verner, William B Jones, Craig M. Hill, James W. Janc, Joseph J. Buggy, Ning Ding, John C. Reneau, Michael S. Khodadoust, Youn H. Kim, Ryan A. Wilcox, Richard A. Miller

## Abstract

ITK is a tyrosine kinase expressed predominantly by T lymphocytes. In mice, selective knock-out of the ITK gene produces Th1 skewing of T helper cell differentiation. We synthesized a covalent ITK inhibitor, soquelitinib, that binds ITK with greater than 100-fold selectivity compared to binding to resting lymphocyte kinase (RLK). *In vitro* studies with normal or malignant T cells demonstrated that soquelitinib suppresses Th2 cytokine production preferentially with relative sparing of Th1 cytokines. Soquelitinib inhibits the *in vivo* growth of several syngeneic murine tumors including those that do not express ITK. Treatment with soquelitinib leads to increased tumor infiltration of normal CD8+ cells that possess enhanced T effector function. Soquelitinib inhibited expression of T cell exhaustion markers and was able to restore T effector function to exhausted cells. Pharmacologic selective ITK inhibition may represent a novel approach to cancer immunotherapy.

## INTRODUCTION

ITK is a member of TEC family of kinases that is expressed in T, NK and mast cells^1^. ITK plays an important role in T cell receptor (TCR) signaling as well in the differentiation of thymocytes into mature T cells^1^. Following TCR stimulation, ITK is recruited to the membrane associated SLP 76/LAT adapter complex, where it is phosphorylated and activated by the src family kinase LCK^2^. Activated ITK then phosphorylates phospholipase C (PLC)γ1, leading to the mobilization of Ca2+ and to the activation of growth, survival, and differentiation pathways regulated by mitogen activated protein kinase (MAPK), nuclear factor kappa B (NFκB), SH2 containing protein tyrosine phosphatase-2 (SHP2)^3^, interferon regulatory factor (IRF)4^4^, and GATA-3 binding protein (GATA-3)^5^. Together, these pathways indicate that ITK plays important roles in TCR signaling, T cell proliferation, and differentiation of naïve CD4+ T cells.

ITK-/- knockout mice exhibit defects in T helper 2 (Th2) differentiation, while retaining the ability to differentiate into Th1 cells that secrete interferon gamma (IFNγ)^6^, a phenomenon known as Th1 skewing. Another TEC-family kinase, resting lymphocyte kinase (RLK/TXK), is also expressed in T cells and is similarly activated by TCR-driven phosphorylation by src-family kinases ^2^. RLK interacts with many of the same signaling components as ITK.^7, 8^ Whereas ITK^-/-^ CD4+ T cells in mice have impaired T cell activation and differentiation, ITK^-/-^ RLK^-/-^ double knockout T cells have a more substantial signaling defect, resulting in a profound loss of normal T cell function^9^. Thus, selective inhibition of ITK while sparing RLK should result in modulation of T cell responses without a marked effect on overall T-dependent immunity.

We hypothesized that selective pharmacologic inhibition of ITK with sparing of RLK would affect T cell differentiation leading to Th1 skewing, which could enhance anti-tumor immunity. Therefore, we have synthesized a novel ITK selective, orally bioavailable ITK inhibitor, soquelitinib (known previously as CPI-818), that binds covalently to cysteine 442 of ITK and blocks its function.

Here we report the biologic effects of selective pharmacologic blockade of ITK. Our findings demonstrate effects on T cell differentiation including Th1 skewing, blockade of Th2 function and reduction and reversal of T cell exhaustion*. In vivo* studies in syngeneic tumor models reveal inhibition of tumor growth that is attributable to soquelitinib -mediated immune modulatory effects. These findings indicate that ITK is a potential novel target to enhance the immunotherapy of cancer.

## METHODS

### Mass spectrometry

ITK was obtained as a glutathione S transferase (GST) fusion protein from Carna Bioscience (Product Number 08 081, Lot 13CBS 0356K). The protein was utilized at 586 µg/mL (5.9 µM ITK) in a buffer containing 50 mM TRIS, 150 mM NaCl, 0.05% Brij35 (Thermo Fisher), 1 mM dithiothreitol, and 10% glycerol at pH 7.5. The protein construct comprised amino acids 2 to 620 of ITK and is fused to GST at the N terminus of ITK (ITK GST). A 2 µM sample of the ITK GST fusion protein was incubated with a 5-fold molar excess of soquelitinib (10 µM). A control, untreated sample of ITK GST was prepared in parallel. The samples were incubated at room temperature for 2.5 hours to allow for complete inhibition of ITK GST by soquelitinib. After the incubation phase, samples were frozen and stored at -80°C. Mass spectral analysis was conducted in collaboration with the Stanford University mass spectrometry facility. The control, untreated ITK GST fusion protein, and the soquelitinib inhibited sample were subjected to chymotrypsin digestion and then analyzed by liquid chromatography coupled with tandem mass spectrometry (LC MS/MS). Peptides were separated using an Acquity M-Class liquid chromatograph (Waters Corporation, Milford, MA). Peptide separation was completed on a silica column packed with 1.9 micron C18 stationary phase. An analytical gradient of 80 minutes was used, with mobile phase A being 0.2% formic acid in water, and mobile phase B being 0.2% formic acid in acetonitirle. Ions electrosprayed from this column were detected using an Orbitrap Fusion mass spectrometer (Thermo Scientific, San Jose, CA) operated in a data dependent fashion using both electron transfer dissociation and high energy dissociation fragmentation to improve detection efficiency of modified peptides. To analyze these data, raw spectral files were investigated using Byonic v2.10.5 (Protein Metrics, Cupertino, CA) in a targeted database for the protein of interest. Peptides were constrained to 12 ppm mass accuracy for precursor ions, and 0.4 Da mass accuracies for fragment ions detected in the ion trap. Peptides were assumed to be digested in a semi specific fashion, and up to 2 missed cleavages were allowed. Assigned observed peptides were then validated by inspection of the initial ion mass spectrum, fragmented ion mass spectrum, and extracted ion chromatogram using Byologic (Protein Metrics, Cupertino, CA).

### Biochemical kinase assays

Two biochemical assays of kinase activity were employed. The first is based on the Invitrogen Lanthascreen platform while the second used the PerkinElmer microfluidic based LabChip 3000 system. In the Lanthascreen assay each kinase (1.0 nM) was incubated with compound for 120 minutes at 22°C in a buffer composed of 50 mM HEPES (pH 7.5), 10 mM MgCl_2_, 1 mM EGTA, and 0.01% Brij 35. The kinase reaction was initiated by the addition of ATP (50 µM final) and fluorescein-poly-Glu-Tyr peptide (200 nM final). The reaction was quenched after 60 minutes at 22°C, and phospho-peptide product detected by the addition of EDTA (10 mM final) and terbium PY20 antibody (2 nM final). The phospho-peptide product was measured by time-resolved fluorescence resonance energy transfer (TR FRET) between the terbium of the antibody and fluorescein of the phospho-peptide using an EnVision plate reader (PerkinElmer, Waltham, MA) equipped to detect the TR-FRET signal (excitation 320 nm, dual emission at 495 nm and 520 nm). All reagents including recombinant human kinases, fluorescein-poly-Glu-Tyr peptide, terbium PY20 antibody, and kinase buffer A were purchased from Invitrogen.

Alternatively, in the microfluidic assay each kinase (0.2 nM) was incubated with compound for 15 minutes at 22°C in a buffer composed of 100 mM HEPES (pH 7.5), 0.1% BSA, 0.01% Triton X-100, 1 mM dithiothreitol (DTT), 5 mM MgCl_2_, 10 µM Sodium Orthovanadate, 10 µM Beta-Glycerophosphate and 1% DMSO. The kinase reaction was initiated by the addition of ATP (10 µM final) and fluorescein labeled Scrtide peptide (AnaSpec, Fremont, CA)(1 µM final). The reaction was quenched after 3 hours at 22°C by the addition of EDTA (20 mM final), and phospho-peptide product detected by direct fluorescence (ex: 480 nm, em: 520 nm) after electrophoretic separation from the substrate using a PerkinElmer EZ reader. All reagents were purchased from Sigma-Aldrich. Inhibition of kinase activity was measured by construction of a concentration response curve of the inhibitor.

### pERK and pS6 flow cytometry

Peripheral blood mononuclear cells (PBMCs) were pelleted (400 x g at 4°C for 5 minutes) and resuspended in RPMI + 10% heat-inactivated fetal bovine serum (FBS) at 14.3 x 10^6^ cells/mL; 70 μL (1 x 10^6^ cells) were plated in a 96 well U bottom plate and incubated at 37°C for 1 hour. A 10 mM stock solution of soquelitinib was prepared in DMSO and then serially diluted to 100, 10, 1, and 0.1 μM; soquelitinib in a volume of 10 μL was added to the cells and incubated at 37°C for 1 hour. Anti-CD3-biotin and anti-CD28-biotin were prepared together at 10 μg/mL each; 10 μL antibodies were added to the cells and incubated at 37°C for 15 minutes. Avidin was prepared at 500 μg/mL; an aliquot of 10 μL avidin was added to the cells and incubated at 37°C for 5 minutes to induce TCR crosslinking. A total volume of 100 μL cells was transferred to 200 μL warm 2.4% paraformaldehyde (PFA) in deep well blocks. Pipetting was performed using a liquidator 96 to add reagents to the entire plate with uniform timing. Cells were fixed with PFA. Cells were pelleted, and the plate was aspirated leaving approximately 100 μL residual volume. The plate was vortexed for 10 seconds, and 1 mL −20°C methanol was added to each well. For antibody staining, cells were pelleted, washed twice with fluorescent-activated cell sorter (FACS) buffer (phosphate-buffered saline [PBS] + 1% bovine serum albumin [BSA] + 0.1% sodium azide), stained at room temperature for 1 hour, washed twice with FACS buffer, fixed with 1.6% PFA, and acquired on a cytoflex flow cytometer. Inhibition was measured by construction of a concentration response curve of the Log base 10 transformed values of the molar concentration of inhibitor.

Cells were stained with the following antibodies against mouse CD45 (30-F1, Biolegend), CD8 (53-6.7, Biolegend), CD4 (GK1.5, Biolegend), PD-1 (29F.1A12, BD), TIGIT (1G9, BD), LAG3 (C9B7W, Biolegend), Tim3 (RMT3-23, Biolegend), CXCR3 (CXCR3-173, Biolegend), TCRVa2 (B20.1, Biolegend), or against human CD3 (OKT3, Biolegend), CD4 (RPA-T4, Biolegend), CD45RA (HI100, Biolegend), CCR7 (G043H7, Biolegend). Fixable Viability Dye eFluor506, and eFuor780 (eBioscience) or ViaKrome 808 Fixable Viability Dye (Beckman Coulter) were used to exclude dead cells before analysis. For intracellular staining, cells were first stained with surface markers and then fix/ permeabilized using the Foxp3/Transcription Factor Staining kit (eBioscience) according to the manufacture’s protocol. For detection of intracellular IFNγ, TNF, cells were stimulated *in vitro* with either PMA (50 ng/ml, Thermo Fisher) and ionomycin (1 μM, Thermo Fisher) or with 5 ug/ml OVA peptide (a.a.257-264, InvivoGen) for 4-5 hr in the presence of Golgi Stop (1/1000, BD) and Golgi Plug (1/1000, BD). Cells were then harvested for surface marker staining, followed by fixation and permeabilization as described above. Permeabilized cells were stained with antibodies against IFNγ (XMG1.2, Biolegend), and TNF (MP6-XT22, Biolegend). For detection of Granzyme B, perforin, and T cell factor (TCF)-1, cells were stained with surface markers prior to fixation and permeabilization. Permeabilized cells were then stained with antibodies against Granzyme B (GB11, BD(Becton Dickinson)), Perforin (S16009A, Biolegend), TCF-1 (S33-966, BD). Flow cytometry was then performed on the Cytoflex LX flow cytometer (Beckman Coulter) and analyzed using Flowjo software (TreeStar).

### Inhibition of IL-2 production

Jurkat cells (0.25 X 10^6^) in Dulbecco’s Modified Eagle’s medium (Gibco) supplemented with 5% FBS were seeded in each well of a 96 well MultiScreen^®^ HV filter plate (EMD Millipore) placed on top of a 96 well collection plate. Soquelitinib was serially diluted in DMSO and added to the culture. After a 2-hour incubation at 37°C the plates were centrifuged (188X *g* for 15 seconds) to remove the inhibitor-containing medium from the filter plate. Cells were washed twice by resuspending with 200 μL of medium per well and centrifuged. Cells in each well were then resuspended with 1.25 X 10^6^ (50 μL) of Dynabeads CD3 (Life Technologies) and incubated for 18 hours at 37°C. Conditioned media were collected by centrifugation (188X *g* for 1 min). Human IL-2 concentration was measured with the AlphaLISA human IL2 Kit (PerkinElmer) and data collected on an Envision Plate Reader (PerkinElmer). Data were analyzed and graphed using GraphPad Prism version b7.0b. Technical duplicates of the concentration response were collected at each determination.

### Western blot analysis

For phosphorylation of phospholipase C (PLC)γ1 and zeta associated protein (ZAP)70, human H9 cells were pre-treated with soquelitinib for 1 hour at 37°C and then stimulated for 30 seconds with 6 μg biotinylated-anti human CD3 (UCHT1, eBioscience) plus 1.5 μg avidin (Pierce). For detection of NF-κB expression, T8ML-1 cells derived from a patient with refractory peripheral T cell lymphoma were unstimulated or stimulated with anti-CD3/28 Dynabeads (Invitrogen) in the presence of soquelitinib or DMSO for one hour. For detection of GATA-3 expression, T8ML-1 cells were left unstimulated or pre-activated with anti-CD3/28 Dynabeads for 24 hours, followed by treatment with either soquelitinib or DMSO for another 24 hours. Nuclear and cytoplasmic extracts from cell lysates were prepared for Western blot analysis as described by Jiang and colleagues^10^. Primary antibodies against total PLCγ1, phosphorylated PLCγ1(Y783), total ZAP70, phosphorylated ZAP70 (Y319), β-actin, ITK, NF-κB (p65), GATA-3, LaminB2 antibodies were purchased from Cell Signaling Technology. IRDye 800CW Anti-Rabbit IgG and IRDye 700CW Anti-Mouse IgG (1:20,000 dilution) secondary antibodies were purchased from LI-COR. Protein signals were detected by the LI-COR Odyssey^®^Fc ImagingSystem.

### Soquelitinib kinase binding activity

The binding constant of soquelitinib to selective kinases including ITK was determined commercially by DiscoverX Corporation (Fremont, CA) using competition binding assays^11^.

### *In vitro* T helper (Th) cell differentiation and human T cell activation

Human PBMCs from healthy donors were isolated by density gradient method using Ficoll-Paque and Sepmate tubes (Stemcell Technology). Live naïve CD4+ T cells were purified from PBMCs post staining with fluorescence labelled antibodies to human CD3(OKT3, Biolegend), CD4 (RPA-T4, Biolegend), CD45RA (HI100, Biolegend), CCR7 (G043H7, Biolegend) followed by sorting of cells using the SONY SH800 sorter or obtained with the naïve human CD4+ T cell isolation kit (StemCell Technology). For T helper cell culture, purified CD4+ T cells were stimulated with immobilized anti-human CD3 (5 μg/ml, clone UCHT1) and soluble anti-human CD28 (1 μg/ml, clone CD28.2) in the presence of Th polarizing cocktails for 6 days (see below). Cells were washed twice with fresh medium and 100,000 differentiated CD4 Th cells were aliquoted to each well and re-stimulated with 12μl of solution mixture of antibodies to human CD2, CD3, and CD28 (ImmunoCult human T cell activator, StemCell Technologies) in total volume of 200 μl complete RPMI medium in a 96-well U bottom plate in the presence of either DMSO or soquelitinib for 48 hours. Culture supernatants were collected, and the amounts of cytokines were assessed by MSD (Meso Scale Discovery cytokine panel). Th1 condition: Recombinant human IFNγ (10 ng/ml, R&D Systems), and IL-12 (10 ng/ml, R&D Systems), anti-human IL-4 (OA19A66, Biolegend) and anti-human IL-5 (TRFK5, Biolegend)neutralizing antibodies (10 ug/ml each, Biolegend); Th2 condition: Recombinant human IL-4 (10 ng/ml, R&D Systems), IL-5 (10 ng/ml, R&D Systems), anti-human IFNγ (B27, BD) and anti-human IL-12 (C8.6, Biolegend) neutralizing antibodies (10 mg/ml each, Biolegend). For T cell activation in non-polarizing conditions, total PBMCs or purified human naïve CD4+ T cells were stimulated with the ImmunoCult human T cell activator (StemCell Technologies) in the presence of either DMSO or soquelitinib for 72 hours. Cytokines were quantified by MSD from culture supernatants or by intracellular cytokine staining. Cells from patients with Sezary syndrome were obtained from patients who were participating in a Phase I trial of soquelitinib to treat T cell lymphoma. Both the treatment protocol and the study of blood from subjects in this trial were approved by the Stanford Medical Center Institutional Review Board.

### Rodent studies

8-10 week old C57BL/6 and Balb/c mice were purchased from Charles River Laboratory. OT-1 TCR transgenic mice were purchased from Jackson Laboratory (stock no. 003831). Safety of orally administered soquelitinib was evaluated in a GLP (Good Laboratory Practice) 28-day repeat dose, toxicity study in Sprague-Dawley rats purchased from Charles River. All procedures involving animals and their care were approved by the Institutional Animal Care and Use Committee and conducted according to institutional guidelines and approved animal protocol.

### Syngeneic murine tumor models

Murine cancer cell lines such as CT26 (colorectal carcinoma), RENCA (renal cell carcinoma), EL4 (T cell lymphoma), A20 (B cell lymphoma) were obtained from ATCC. Ovalbumin-expressing B16F10, B16F10-OVA (melanoma) was purchased from Millipore-Sigma. All cell lines were maintained under limited passage from original stocks (typically under 5). Cells in PBS were mixed in 1:1 volume with matrigel (Corning) and subcutaneously implanted into the right flanks of mice (unless otherwise stated). For tumor growth studies, tumor volume was measured using the formula as follows: Volume= (width^2^ x length)/2. Mice with established tumors (75-100 mm^3^, approximately 8-10 days post engraftment) were randomized and divided into indicated treatment groups.

### *In vivo* treatments

For soquelitinib treatment, a soquelitinib stock was formulated in solution consisting of Tween 80/Kolliphor ELP/propylene glycol/PEG 300/2% hydroxypropyl cellulose water or in chow diets with macronutrient levels matching the common Teklad 2018 diet (Research Diets, Inc.). Tumor-bearing animals were treated with solution-formulated soquelitinib or vehicle control by oral gavage twice a day (30 mg/kg/day), unless otherwise indicated in figure legends or allowed access to *ad libitum* control or soquelitinib diets (130 or 200 mg/kg/day). Adequate drug exposure was confirmed by standard LC-MS method in plasma samples. For immune checkpoint blockade (ICB), suboptimal dose (25 μg /mouse) of isotype antibodies (rat IgG2a, or Syrian hamster IgG, BioXcell) or anti-PD-1 (RPM 1-14, BioXcell) and/or anti-CTLA4 (9H10, BioXcell) blocking antibodies were injected every three days intraperitoneally. For*in vivo* antibody depletion, anti-CD4 (GK1.5, 200 μg /mouse), anti-CD8 (53-6.72, 200 μg /mouse), anti-NK1.1 (PK136, 200 μg/mouse) antibodies were injected intraperitoneally one day before soquelitinib monotherapy and every three days following the first injection. Cell depletion was verified by flow cytometry using mouse splenocytes.

### Isolation of tumor infiltrating lymphocytes (TILs)

Tumors were collected and digested for 30 minutes at 37°C with the mouse tumor dissociation kit (Miltenyi) in combination with mechanical force using the gentleMACS Octo Dissociator (Miltenyi). Single-cell suspensions were further prepared as previously described^12^ and then used in various assays of T cell function.

### CD8 Immunohistochemistry (IHC)

The assessment of CD8+ tumor infiltrating lymphocytes (TILs) was performed by immunohistochemistry (IHC) on sections from formalin-fixed, paraffin-embedded EL4 tumor samples using the Ventana Discovery Ultra Platform. Sections were staining with anti-mouse CD8 antibody (EPR21769, Abcam) together with the Ventana Discovery ChromoMap Kit, followed by the secondary rabbit IgG antibody. Visualization was obtained with the Ventana Discovery OmniMap Anti-Rabbit HRP kit. Indica Labs Halo Software CytoNuclear 2.0.5 Algorithm was used for image analysis.

### Analysis of transcripts associated with exhaustion by Nanostring

CD8+ tumor infiltrating lymphocytes (TILs) were enriched using the mouse CD8 TIL enrichment kit (Miltenyi) before total RNA extraction. RNA sample preparation and Nanostring analysis were performed as previously described^13^. Assessment of expression of exhaustion genes were done using the Nanostring mouse Pancancer IO360 panel and the data were normalized using Nanostring nSolver software. Z-score transformed data were then analyzed to determine significant changes in gene expression among different treatment groups. For each gene set, mean from 3 tumor samples per treatment group was compared and displayed as fold changes to the vehicle-treatment group.

### *In vitro* T cell exhaustion culture

Pooled spleen and lymph nodes from the OT-1 TCR transgenic mice were used to generate single-cell suspensions after RBC lysis. OVA peptide (a.a.257-264, InvivoGen) was added at concentration of 5 μg/ml to washed pooled cells and cultured as 5 x 10^6^ cells/ml. After 48 hours, co-cultures were counted and recultured as 1 x 10^6^ cells/ml in fresh medium containing 20 U/ml recombinant human IL-2 (StemCell Technologies) in the presence of soquelitinib or DMSO for additional 5 days. Every 2 days, OVA peptide was added for repeated stimulation and fresh soquelitinib was replenished in the culture. On day 7, cells were either harvested to assess levels of exhaustion marker expression by flow cytometry or DMSO-treated cells were purified using the mouse CD8 T cell isolation kit (StemCell Technologies). Enriched mouse CD8+ T cells were then treated with either DMSO alone or soquelitinib at concentrations ranging from 0 to 10 μM with additional two rounds of stimulation with the OVA peptide. Cells were harvested on day 11 to measure granzyme B and IFNγ cytokine production. For *in vitro* exhaustion culture using human CD4 T cells, naïve CD4 T cells were purified using the naïve human CD4+ T cell isolation kit (StemCell Technology) and stimulated with immobilized anti-human CD3 (5 μg/ml, clone UCHT1) and soluble anti-human CD28 (1 μg/ml, clone CD28.2) for 48hr without soquelitinib. Stimulated CD4+ T cells were passaged onto a fresh anti-CD3 coated plate every two days for additional 3 times in the presence of 5 ng/ml hIL-2 and soquelitinib at the indicated concentration. On day 8, cells were harvested for the assessment of exhaustion marker expression.

### Statistical analysis

The data are displayed as means ± SEM, unless otherwise indicated. All statistical analyses were performed using GraphPad Prism Software. Significance was determined using an unpaired two-tailed Student t-test, unless otherwise indicated. Differences were considered statistically significant at p<0.05. P values are reported as: ns. p> 0.1, *p <0.05, **p <0.01, ***p <0.001, ****p <0.0001.

## RESULTS

### Chemical Design and Synthesis

Analysis of ligand bound ITK crystal structures in the Protein Data Bank (PDB) suggested the aminothiazole-based molecular scaffold (PDB entry 3MJ2) as a suitable starting point for structure-based drug design of ITK-selective covalent inhibitors. This ligand forms two hydrogen bonds with the Met-438 residue in the hinge region and a hydrophobic interaction with the gatekeeper residue Phe-435. A piperazine-acetamide fragment projects toward Cys-442, providing a useful vector for ligand modification with electrophilic substituents to engage Cys-442 in covalent bond formation. The benzylamine moiety was replaced with the less bulky and less polar cyclopropyl-substituent. Further optimization of ITK selectivity and potency was achieved with the introduction of the 7-membered 1,4-diazepane ring with the R-methyl group. Presumably, the chiral methyl acts as an anchor to stabilize one of the conformations of the 7-membered ring that directs the acrylamide fragment towards Cys-442 facilitating irreversible covalent bond formation between soquelitinib and ITK via a Michael addition reaction. The chemical structure of soquelitinib (M.W.= 514.66) and selective binding are shown in Figure 1, Panels A and C.

**Figure 1.**
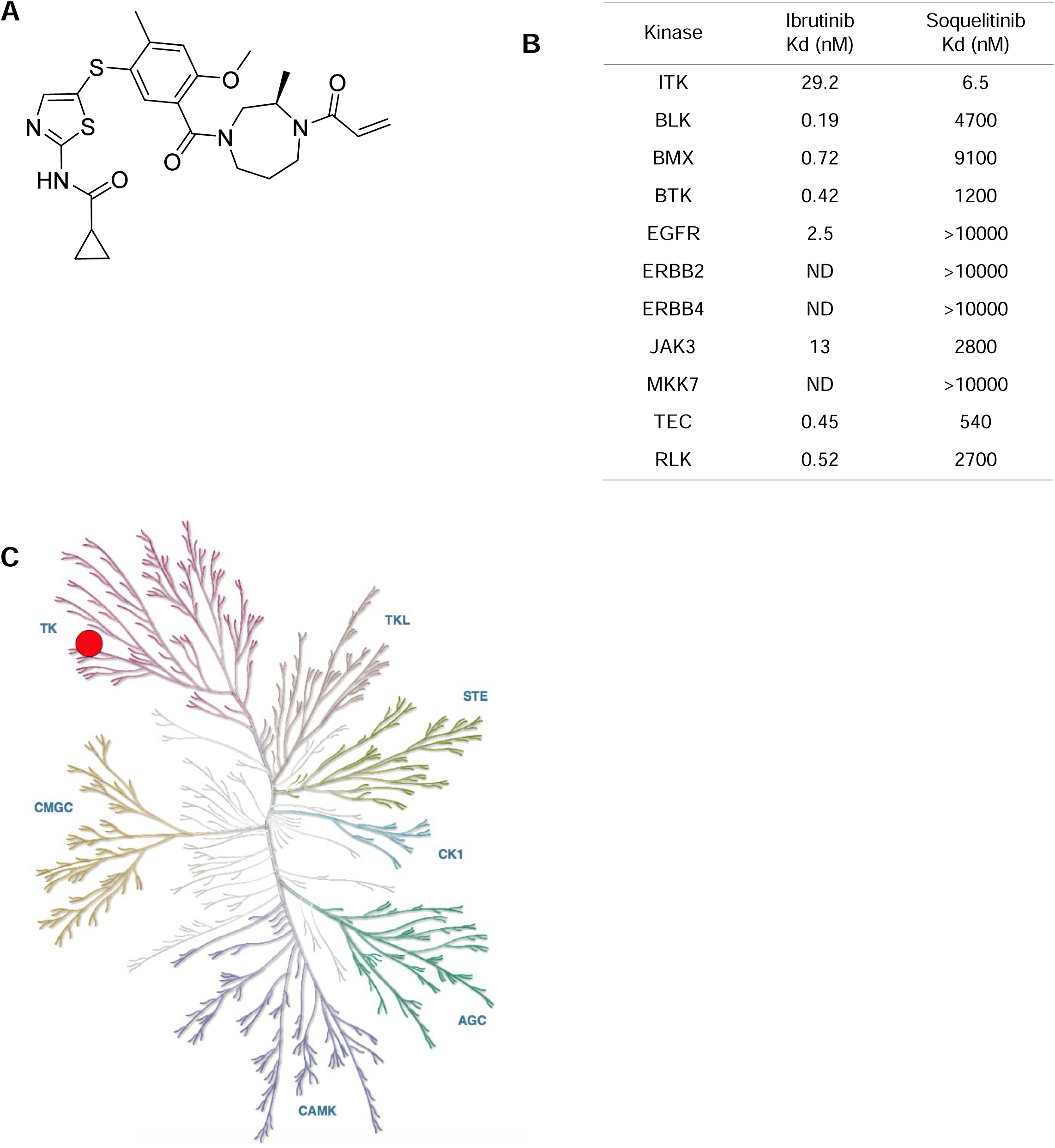
Structure and specificity of soquelitinib. (A) Chemical structure of soquelitinib. (B) Eleven kinases with a cysteine located comparably to the cysteine at amino acid 442 of ITK were studied. The specificity of soquelitinib binding compared to the specificity of ibrutinib binding. (C) The specificity of soquelitinib is represented as a kinome map. Abbreviations: TK=tyrosine kinase; TKL=tyrosine kinase like; STE, AGC, CMGC are families of kinases; CK1=casein kinase 1; CAMK=calcium/calmodulin dependent protein kinase; BLK=B lymphocyte kinase; BMX=a non-receptor protein kinase; BTK=Bruton’s tyrosine kinase; EGFR=epidermal growth factor receptor; ERBB2 and ERBB4=epidermal growth factor receptor family members; JAK3=Janus kinase 3; MKK7=MAP kinase kinase 7; TEC=a member of a family of non-receptor tyrosine kinases; RLK=resting lymphocyte kinase; ND=not determined.

### Covalent irreversible inhibition of ITK by soquelitinib

A chymotryptic digest of soquelitinib -inhibited ITK was analyzed by liquid chromatography with tandem mass spectroscopy (LC MS/MS) to detect the presence of the covalently linked inhibitor to the enzyme. The digest generated 247 peptide fragments and their sequences were identified through deconvolution of the mass spectra. A set of three nested peptides were detected in the sample treated with soquelitinib that were absent from the untreated control sample (Supplemental Table 1). Each of the three peptides contained Cys-442 and their masses were shifted by the mass of soquelitinib when compared to the corresponding unmodified peptide sequence. The presence of the three unique peptide fragments generated by soquelitinib treatment is consistent with the inhibitor covalently labeling Cys-442. There were no other peptide fragments detected that were modified by soquelitinib. Collectively these data demonstrate that soquelitinib is a selective, irreversible, and covalent inhibitor of ITK.

### Kinase selectivity of soquelitinib

The five members of the TEC kinase family and six other enzymes of the human kinome contain a cysteine in a position homologous to Cys-442 in ITK and could potentially be irreversibly inhibited by soquelitinib ^14^. To investigate the selectivity against the 11 cysteine-containing kinases, Kd values for soquelitinib for each enzyme were obtained. Soquelitinib displayed aKd = 6.5 nM for ITK and was at least 80-fold selective over the remaining cysteine-containing kinases (Figure 1B). Profiling data for ibrutinib, a covalent BTK inhibitor, approved for treatment of B cell lymphomas also is shown for comparison. Soquelitinib was then profiled against the human kinome at the single concentration of 1.0 µM. Soquelitinib displayed a high degree of selectivity such that only eight kinases were inhibited ≥65% and only the targeted kinase, ITK, was inhibited by >95% (Figure 1C). We measured k_inact_/*K*_i_ values for the two kinases, ITK and RLK, and found soquelitinib is 115-fold selective toward ITK over RLK. Moreover, other kinases in the T cell receptor pathway were not inhibited by soquelitinib (kinase activity, percent control: Lck, 89; Fyn, 99; ZAP-70, 59; RLK, 70).

### Soquelitinib inhibits TCR signaling downstream of ITK and blocks IL-2 production

Figure 2A shows the proposed signaling pathways of ITK. To investgiate whether inhibition of ITK activity by soquelitinib modulates TCR signaling during activation, we measured phosphorylation of several proteins downstream of TCR signaling such as ZAP70, PLCγ1 in human H9 cells, and ERK and ribosomal protein S6 in human CD4+ T cells, respectively. As shown in Figure 2B and 2C, soquelitinib inhibited the phosphorylation of PLCγ1, ERK and S6 protein and had a minimal impact on phosphorylation of a proximal kinase, ZAP70. Previous study from Wilcox’s group^10 15^ has shown that TCR-medicated activation of NF-κB and GATA-3 is ITK dependent. We tested whether blockade of ITK kinase activity by soquelitinib could affect GATA-3 expression which plays an essential role in Th2 cell differentiation. Consistent with the published report, we observed impaired NF-κB activation after soquelitinib treatment for 60 minutes in the T8ML-1 cell line, a cell line derived from a patient with refractory peripheral T cell lymphoma (PTCL) as shown by reduced NF-κB protein levels in both cytoplasmic and nuclear fractions (Figure 2D). We found that soquelitinib treatment lowered the expression of both ITK and GATA-3 as expected since ITK expression is regulated by GATA-3^15^ (Figure 2E). Together, these results indicated that inhibition of ITK kinase activity by soquelitinib led to inhibition of downstream TCR signaling events and GATA-3 expression. We next investigated the effect on IL-2 secretion, an early event in T cell activation. Soquelitinib suppressed IL-2 secretion in Jurkat cells in response to TCR stimulation with a mean IC_50_ of 136 nM (Fig. 2F). Given that IL-2 plays an essential role in T cell proliferation and survival, we evaluated the viability and proliferation of human T cells treated with a range of soquelitinib doses. As shown in Figure 2G, our results revealed that at 10uM concentrations of soquelitinib, cell viability was unaffected but total cell number was reduced, findings consistent with an antiproliferative effect seen only at the highest concentration. Viability and proliferation were not affected at lower concentrations of drug.

**Figure 2.**
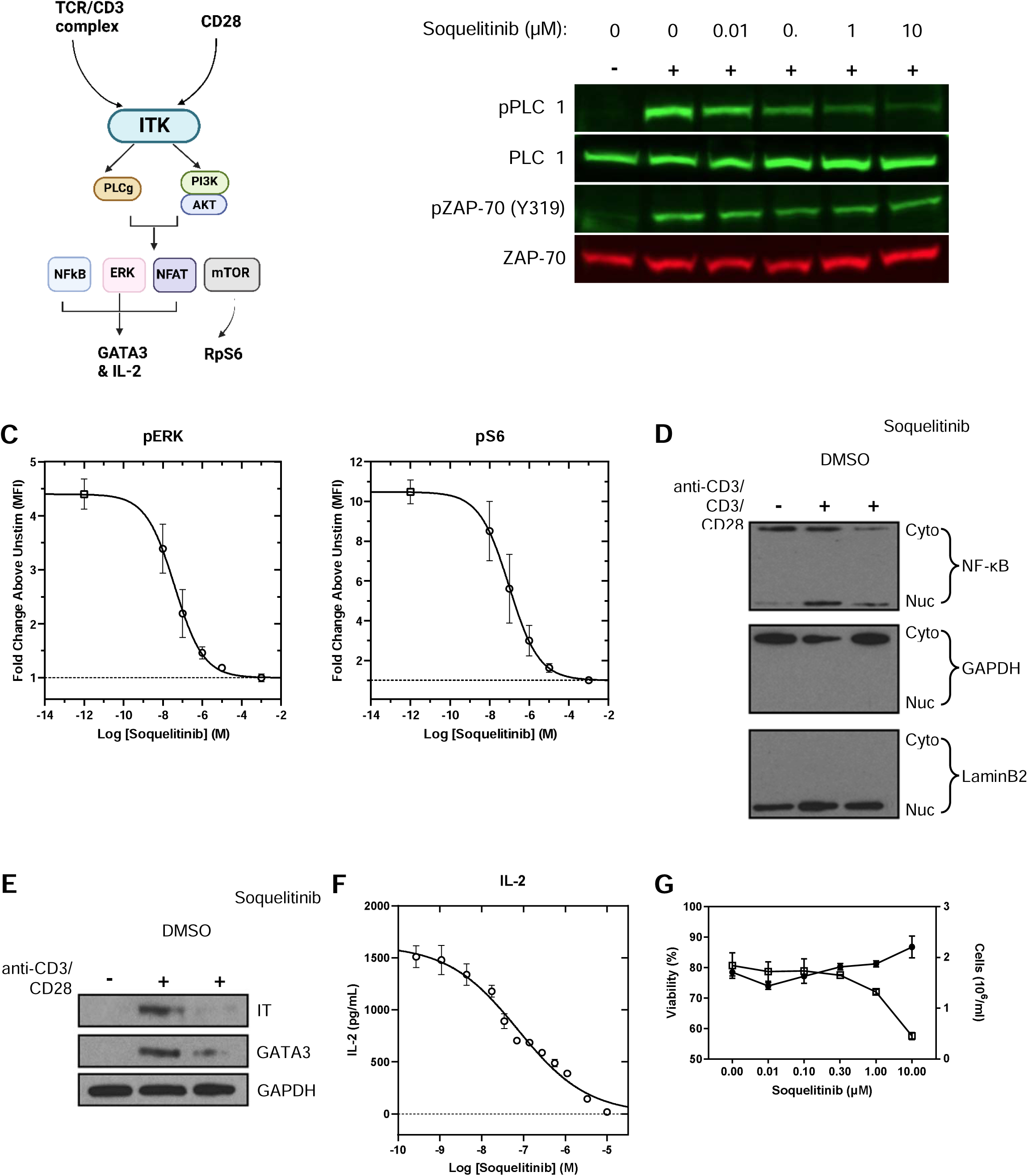
Soquelitinib inhibits TCR-induced downstream signaling and activation. (A) Diagram illustrates pathways that activate ITK and the downstream effects of that activation. (B) Western blot analysis of whole-cell lysates from H9 cells stimulated with anti-CD3 for 30 seconds in the presence of increasing concentration of soquelitinib. Blots were probed with specific antibodies to total and phosphorylated PLCγ1 and ZAP-70. (C) soquelitinib inhibits ERK and S6 phosphorylation. ERK and S6 phosphorylation was measured by flow cytometry following anti-CD3/28-crosslinking in human T cells. Each data point displays mean of fold change from three donors. Error bars represent SEM. (D) Reduced TCR-induced nuclear translocation of NF-kB following soquelitinib treatment. Cytoplasmic (Cyt) and nuclear (Nuc) immunoblot analysis of T8ML-1 cells that were unstimulated or stimulated with anti-CD3/CD28 beads for 60 min in the presence of either DMSO or 10 mM of soquelitinib. Blots were probed with anti-NF-kB (p65), GAPDH, and LaminB2. (E) Decreased GATA3 expression following soquelitinib treatment. Western blot analysis of whole-cell lysates derived from T8ML-1 cells that were unstimulated or activated 24 hr prior to whole day treatment with soquelitinib (10 mM) or DMSO. (F) Soquelitinib suppresses IL-2 secretion in Jurkat cells in response to TCR stimulation. A representative curve is shown from one experiment with the mean of technical duplicates plotted. (G) Soquelitinib (≥ 1 µM) reduced the total number of cells while maintaining cell viability. Cell viability and cell expansion rate of human primary T cells measured by the Vi-Cell XL Cell Viability Analyzer (Beckman Coulter). Data are means ±SEM of technical triplicates within an experiment and are representative of 15 human donors. Abbreviations: PI3K=phosphatidyl inositol-3 kinase; AKT=protein kinase B; NFκB=nuclear factor kappa B; NFAT=nuclear factor of activated T cells; ERK=extracellular signal regulated kinase; mTOR=molecular target of rapamycin; GATA3=GATA3 binding protein; IL=interleukin; RpS6=ribosomal protein 6.

### Soquelitinib suppresses Th2 associated cytokines and induces Th1 skewing in CD4 T cells

We evaluated the effect of soquelitinib on T effector cytokines following 3 days of TCR stimulation in purified CD4 T cells from either healthy donors (Figure 3A) or Sezary cells from patients with cutaneous T cell lymphomas (CTCL), which highly express GATA-3 and have a Th2 phenotype ^5, 15^. When normal human T cells are incubated with increasing concentrations of soquelitinib, Th2 cytokine production (IL-4, IL-5 and IL-13) is reduced. The prototypical Th1 cytokine, IFNγ, is affected but only at the highest concentrations of soquelitinib which were tested. Interestingly, Sezary cells exhibited greater sensitivity to soquelitinib as soquelitinib inhibited production of Th2 associated cytokines to a greater extent in Sezary cells than in normal CD4 T cells. Again, their production of IFNγ can also be reduced, but only at the highest concentration of soquelitinib.

**Figure 3.**
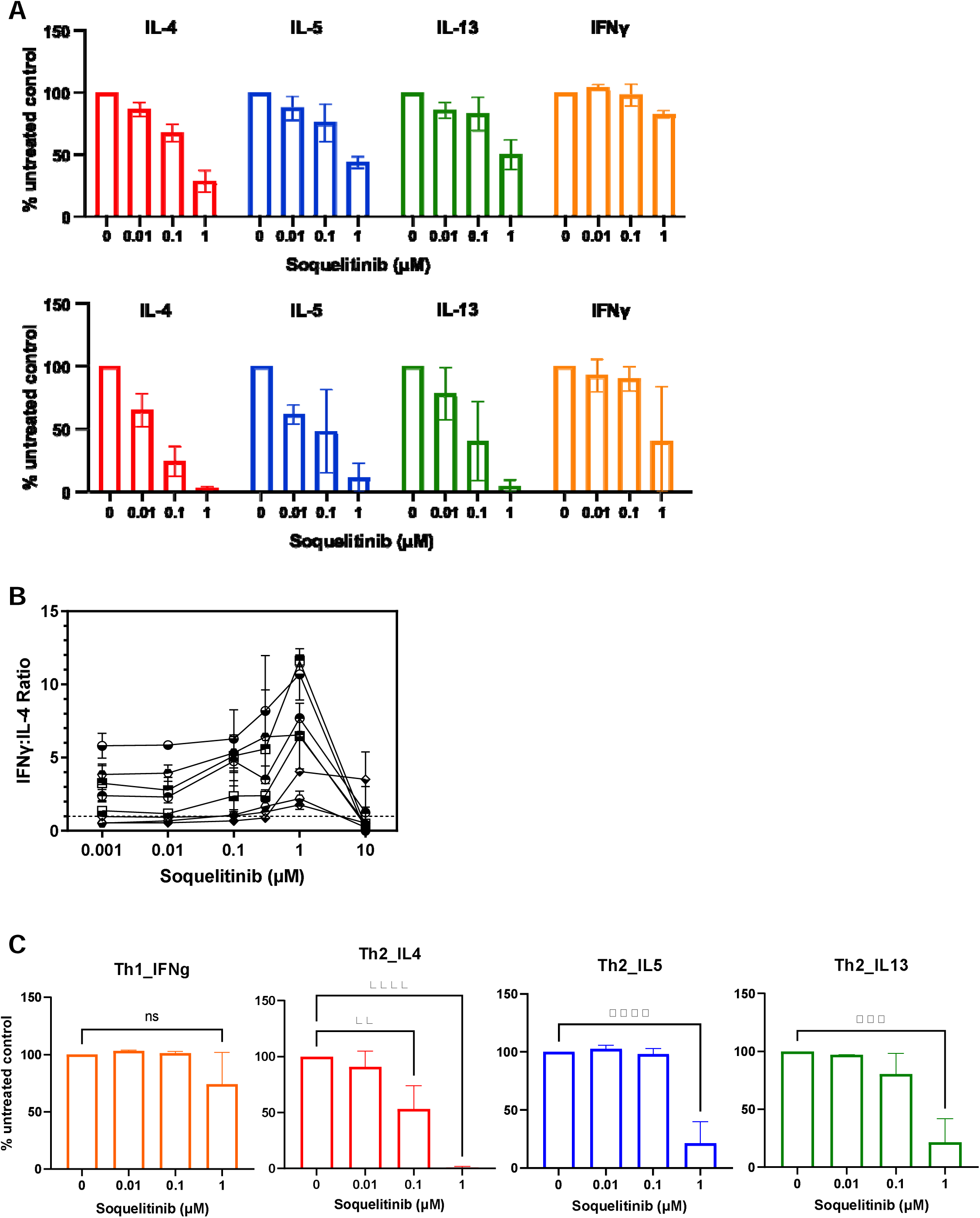
Selective ITK inhibition by soquelitinib skews human T cells toward the Th-1 phenotype. (A) Soquelitinib reduces TH2 effector cytokines in CD4 T cells from either healthy donors (n=3, top panel) or Sezary cells from cutaneous T cell lymphoma (CTCL) patients (n=2, bottom panel). Sorted CD4+ T cells from the blood of healthy donors and the CTCL patients and were stimulated with the ImmunoCult human T cell activator for 3 days in the presence of soquelitinib or DMSO (dimethylsulphoxide) alone. Cytokines in culture supernatants collected on day 3 were quantified by MSD (Meso Scale Discovery cytokine panel). Data were reported as relative to DMSO-treated samples. (B) Soquelitinib (1 μM) induced Th1 skewing with a 2-fold increase in the ratio of IFN(interferon)γ to IL-4. Human PBMCs(peripheral blood mononuclear cells) from 12 donors were pretreated with varying concentrations of soquelitinib for 1hr, followed by TCR (T cell receptor) stimulation. Stimulated cells were cultured for 6 days before restimulation with PMA(phorbol myristate acetate) and ionomycin for 4 additional hours. IFNg and IL-4 production in human CD4 T cells was measured by intracellular cytokine staining. (C) Polarized Th2 but not Th1 CD4+ cells showed greater sensitivity to soquelitinib. Naive CD4+ cells from three donors were first polarized in either Th1 or Th2 polarizing conditions for 6 days and then restimulated by anti-CD3/CD28 in the presence of varying concentrations of soquelitinib or DMSO for additional 2 days. Cytokines in culture supernatants collected after 48 hrs treatment and quantified by MSD. NS, not significant, * p ≤ 0.05, ** p ≤ 0.005, *** p ≤ 0.001 and **** p ≤ 0.0001. (One-way ANOVA). Data displayed as means ± SEM.

We further validated the effect of soquelitinib on Th1 skewing of naïve CD4 T cells by enumerating the intracellular ratio of IFNγ to IL-4. Naive human CD4 T cells were activated in the presence of varying concentrations of soquelitinib for 6 days (Figure 3B). Soquelitinib had a minimal effect on the ratio of IFNγ+/CD4+ T cells at doses up to 1 μM. In contrast, the percentage of IL-4+CD4+ T cells was significantly reduced at 0.3 and 1μM. As a result, at 1 μM concentration of soquelitinib, the ratio of IFNγ to IL-4 CD4+ T cellswas increased by at least 2-fold (Figure 3B). At a concentration of 10μM, soquelitinib completely abolished production of either IL-4 or IFNγ in naïve CD4+ T cells, possibly due to its antiproliferative effects at high concentrations. Together, our results indicated that soquelitinib induced a Th1-skewing phenotype in naïve CD4 T cells in a non-polarizing culture condition.

Next, we evaluated the effect of ITK inhibition on already polarized CD4 helper cells. Naïve CD4 cells were stimulated and differentiated under Th1 orTh2 polarizing conditions for 6 days before soquelitinib was introduced into the culture conditions for an additional three days. Similar to ITK inhibition in naïve CD4 T cells, we found that soquelitinib inhibited the synthesis of Th2-associated cytokines such as IL-4, IL-5 and IL-13 in already polarized Th2 cells, while IFNγ production by Th1 was minimally affected (Figure 3C). Collectively, our results indicated that ITK inhibition by soquelitinib can skew the balance between Th1 and Th2 cells.

### Safety study

The safety of soquelitinib was evaluated in rats given 28 daily oral doses and revealed a no adverse effect level exceeding 1000mg/kg.

### Soquelitinib shows efficacy as single agent in several murine tumor models

Soquelitinib is orally bioavailable and exhibits sustained ITK target occupancy*in vivo*(Supplementary Figure 1). These data prompted us to test the potential of soquelitinib as an immunotherapy for cancer. We first investigated the activity of soquelitinib on tumor growth on a range of murine syngeneic tumor models. These tumor models included CT26 colon adenocarcinoma, RENCA renal cell carcinoma, B16F10 OVA melanoma, A20 B cell lymphoma and EL4, an ITK expressing T cell lymphoma. As shown in Figure 4A, administration of soquelitinib as single agent at 30 mg/kg in solution formulation or 130 mg/kg in chow formulation (A20 model) resulted in a significant inhibition of tumor growth in all five tumor models.

**Figure 4.**
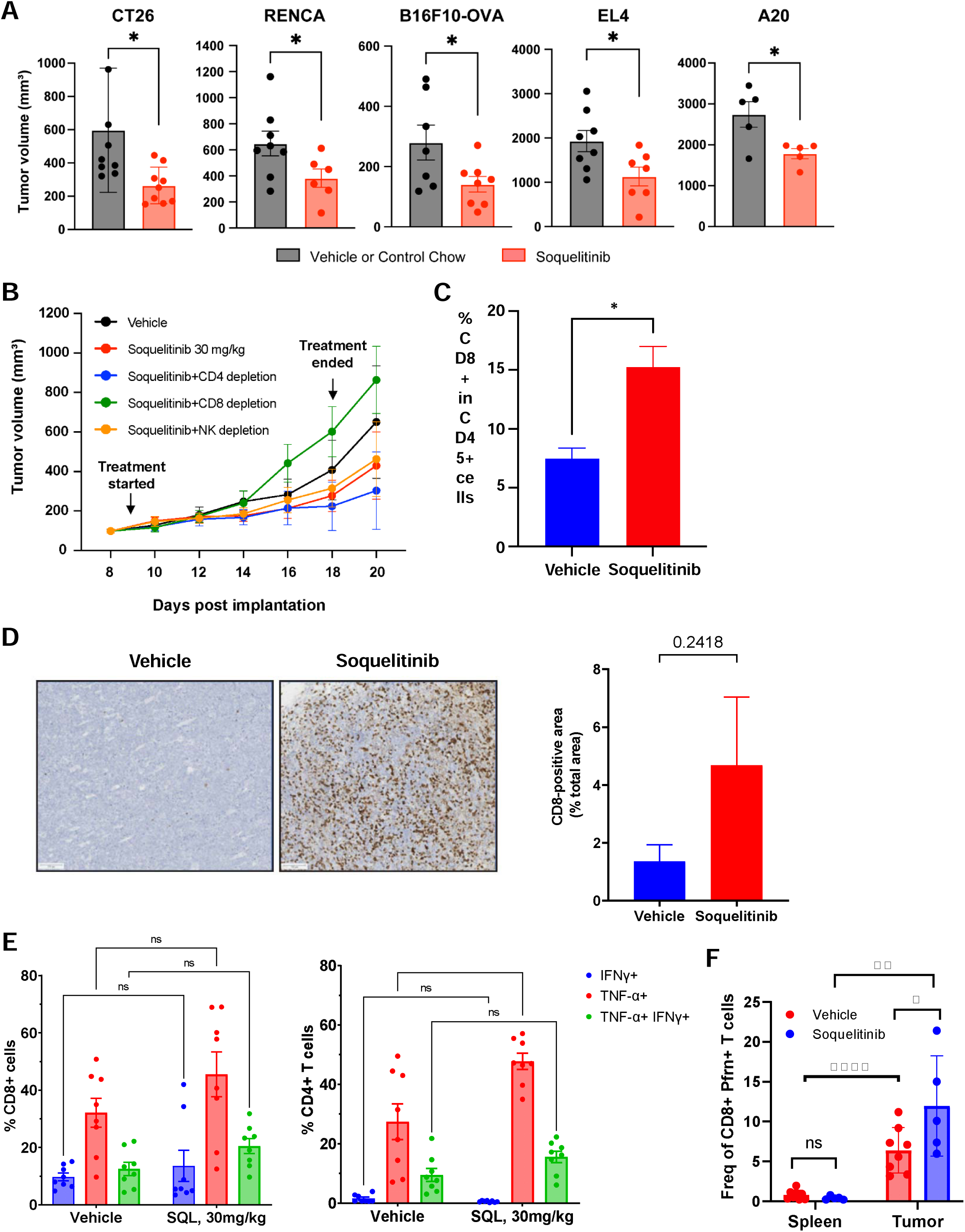
Soquelitinib inhibits tumor growth and enhances cytolytic capacity of tumor infiltrating T cells. (A) Soquelitinib elicits anti-tumor activity in a range of murine syngeneic tumor models. Mice with established tumors indicated here were treated with vehicle/control chow or solution-formulated soquelitinib, 30 mg/kg (CT26 and EL4), 10 mg/kg (RENCA and B16F10-OVA) or soquelitinib chow, 130 mg/kg (A20) daily for 7-8 days (all tumors except A20) or for 13 days (A20). Tumor measurements were performed 2-3 days after last dosing. Results are representative of at least two independent experiments. n=5-8/group. (B) Depletion of CD8+ T cells abolishes antitumor efficacy in the CT26 tumor model. Antibody depletion started one day prior to vehicle or soquelitinib (30mg/kg) treatment. Mice (n=8/group) were treated via oral gavage with solution-formulated soquelitinib or vehicle control for 9 days. Results are representative of two independent experiments. (C) Soquelitinib treatment Increases intratumoral CD8 infiltration in the CT26 tumor model. Percentage of CD8+ TILs was determined by flow cytometry from isolated CD26 tumors treated with either vehicle or soquelitinib for 8 days. (D) Soquelitinib monotherapy induces a higher frequency of intratumoral CD8 T cells in the EL4 model. Representative CD8 immune-histochemistry (left) and quantification of CD8+ area in whole slide section (right, n=3/group) showing the frequency of CD8+ T cells in the EL4 tumors after 8 day of soquelitinib treatment. (E) Soquelitinib -treated animals showed increased production of inflammatory cytokines in tumor infiltrating lymphocytes (TILs) from the CT26 tumors. IFNg- and/ or tumor necrosis factor (TNF)-producing T cells were assessed by intracellular cytokine staining after restimulation *ex vivo* with PMA+ionomycin for 4 hrs. (n=8 per group) (F) Soquelitinib 818-enhances cytotoxicity of CD8+ TILs (tumor infiltrating lymphocytes) purified from the EL4 tumors. Cytotoxicity was assessed as percentage of perforin+ CD8+ T cells. Each symbol represents an individual mouse tumor or spleen. NS, not significant, * p ≤ 0.05, ** p ≤ 0.005, *** p ≤ 0.001 and **** p ≤ 0.0001. Data displayed as means ± SEM. NK=natural killer cell. SQL=soquelitinib.

### CD8 T cell depletionincreases tumor growth in soquelitinib -treated mice

We then investigated the contribution of cell types to the antitumor effect of CPI-818 by *in vivo* antibody depletion of CD4, CD8 or NK cells in the CT26 tumor model (Figure 4B). As expected, treatment with soquelitinib alone significantly inhibited CT26 tumor growth compared with the vehicle group. CD8 T cell depletion of soquelitinib treated mice markedly reduced the anti-tumor activity of soquelitinib (Figure 4B) indicating that the anti-tumor effects of soquelitinib were predominantly mediated by CD8+ T cells (Figure 4B). Depletion of NK or CD4+ cells had no effect on efficacy in the CT26 model (Figure 4B). Together,our data indicate that normal host CD8+ T cells contributed to soquelitinib mediated antitumor response in the CT26 model.

### Soquelitinib treatment enhances CD8 T cell infiltration and increases cytolytic capacity of tumor-infiltrating lymphocytes (TILs)

Since soquelitinib mediated antitumor response is dependent on CD8 T cells, we next examined the effect of soquelitinib on CD8+ T cell infiltration or activation. As shown in Figure 4C, soquelitinib monotherapy induced significantly higher frequency of intra-tumoral CD8 T cells in the CT26 model. Similar to the CT26 tumor model, we observed a moderate increase in intra-tumoral CD8 infiltration upon soquelitinib treatment compared with the vehicle control as quantified by CD8 immunohistochemistry in EL4 tumor samples (Figure 4D). We further assessed cytolytic capacity of infiltrating T cells by measuring inflammatory cytokine production, and perforin production by CD8+ T cells (Figure 4E & F). We demonstrated that soquelitinib alone promoted production of IFNγ and TNF by both CD8+ and CD4+ tumor infiltrating lymphocytes (TIL) within CT26 tumors, most notably in TNF production (Figure 4E). Consistent with the findings in the CT26 model, we also observed an increase in perforin-producing CD8+ T cells obtained from EL4 tumors but not from uninvolved spleens from mice treated with soquelitinib. (Figure 4F). Collectively, our results revealed that soquelitinib enhanced infiltration of TILs in both tumor models, and effector functions of cytotoxic T cells are not impaired by soquelitinib.

### Soquelitinib enhances anti-PD1 and anti-CTLA4 efficacy in the CT26 tumor model

Given that soquelitinib showed activity as single agent in several murine tumor models, we next asked whether soquelitinib could enhance the antitumor activity of anti-PD-1 and anti-CTLA4 antibodies that target Immune checkpoints. As shown in Figure 5A, syngeneic mice with established tumors were engrafted with CT26 tumor cells and randomized into 4 arms (vehicle, soquelitinib alone, dual combination therapy with suboptimal dose of anti-PD1 and anti-CTLA4 antibodies or triple combination therapy with soquelitinib). While soquelitinib or dual combination therapy with anti-PD1 and anti-CTLA4 resulted in substantial inhibition of tumor growth, combining soquelitinib with anti-PD1 and anti-CTLA4 further reduced tumor growth. By day 34 (18 days off treatment), complete tumor regression was identified in 19 out of 20 (95%) mice treated with triple combination therapy from three independent studies. Together, soquelitinib augments the antitumor efficacy of immune checkpoint blockade (ICB), leading to durable antitumor response.

**Figure 5.**
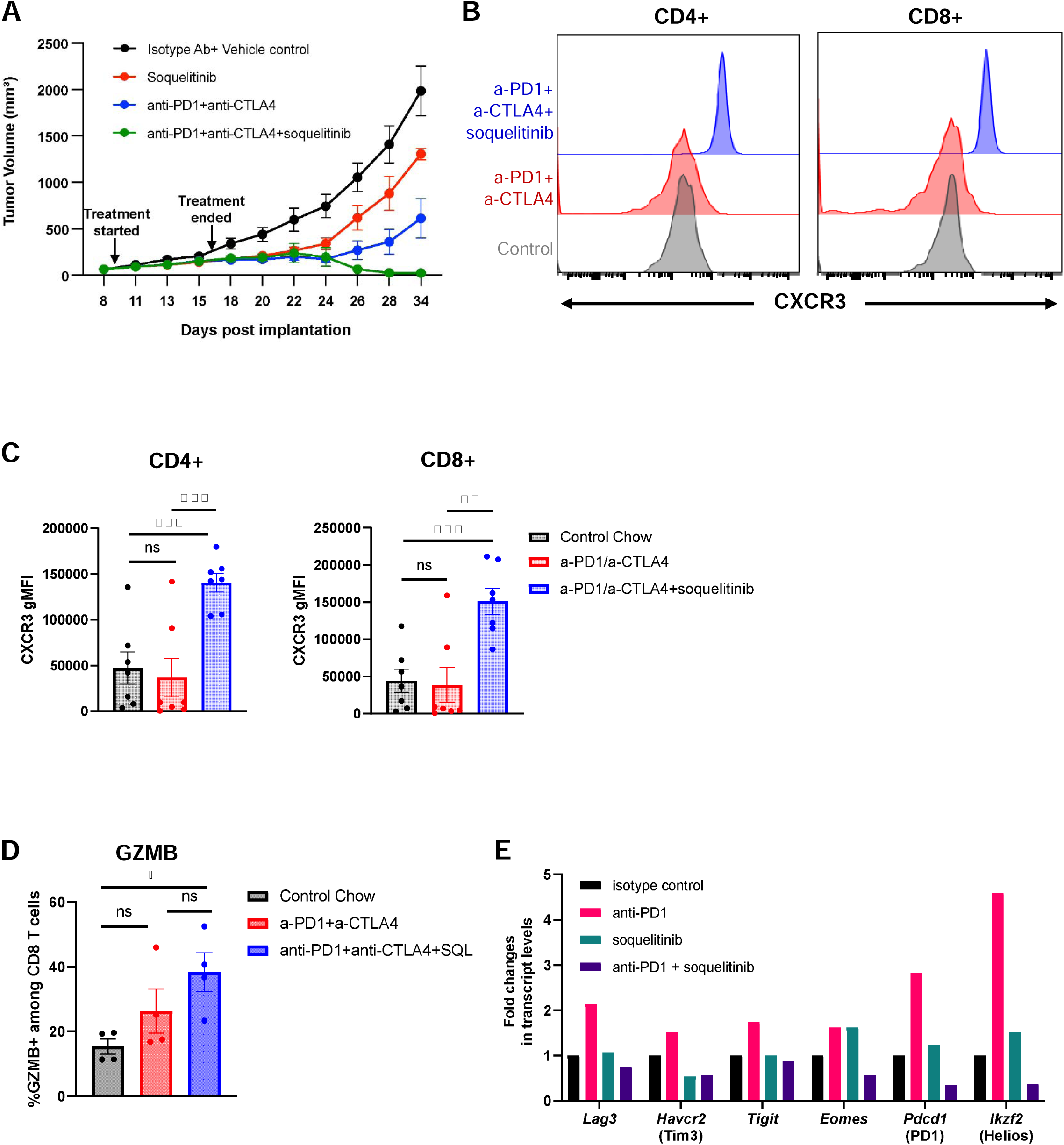
Soquelitinib synergizes with immune checkpoint blockade therapy. (A) Soquelitinib in combination with anti-programming death receptor one (PD1) and anti-cytotoxic T lymphocyte antigen (CTLA)4 induces long-term anti-tumor immunity in the CT26 tumor model. Treatment with solution-formulated soquelitinib alone (30mg/kg) or suboptimal dose of anti-PD1 (25 mg/mouse) and anti-CTLA4 (25 mg/mouse) resulted in an incomplete inhibition of tumor growth, whereas triple combination therapy exhibited durable anti-tumor response even after treatment was terminated for more than 2 weeks. n=8 per group. (B &C) Soquelitinib treated T cells in blood display an increase in CXCR3 cell surface expression. Representative histogram overlays of CXCR3 expression on CD4+ and CD8+ T cells from blood samples (B) and summary graphs of CXCR3 geometric mean fluorescence intensity (gMFI) in CD4+ and CD8+ T cells (C). n=7-8 per group (D) Soquelitinib enhances Granzyme (GZMB) B production in CD8+ TILs. n=4 per group. Blood samples (B &C) or tumors (D) were collected from the CT26 tumor-bearing animals treated with either control chow, anti-PD1+anti-CTLA4 or triple combination therapy with soquelitinib chow (200 mg/kg) for 10 days. (E) Soquelitinib in combination with anti-PD1 reduces transcript levels of exhaustion genes in CD8+ TILs. Transcriptional profiling of CD8+ TILs from CT26 tumors of indicated treatment groups (n=3 per group). NS, not significant, * p ≤ 0.05, ** p ≤ 0.005, and *** p ≤ 0.001. Data displayed as means ± SEM. SQL=soquelitinib.

### Combination of soquelitinib with checkpoint inhibitor blockade enhances CXCR3 upregulation in T cells

We next asked whether soquelitinib -induced Th1 skewing phenotype plays a role in the therapeutic synergy between soquelitinib and checkpoint inhibitors. To track CD4+ Th1 cells*in vivo*, we compared CXCR3 expression, a surrogate marker for Th1 cells, in T cells from the blood of control, anti-PD1 and anti-CTLA4 or triple combination therapy groups. We found a marked upregulation of CXCR3 surface expression in both CD4+ and CD8+ T cells in the triple combination therapy group compared with either control or immune checkpoint blockade (ICB) treatment groups (Figure 5B & 5C). Induction of CXCR3^high^-expressing CD4+ T cells supports the notion that combining soquelitinib with ICB enhances Th1 immunity. Surprisingly, soquelitinib treatment also induced higher levels of CXCR3 expression on CD8 T cells (Figure 5B & 5C). It has been reported that CXCR3 expression on CD8 T cells is required for anti-PD1 mediated antitumor response^16^. We assessed whether CXCR3 upregulation on both CD4 and CD8 T cells correlates with enhanced intra-tumoral CD8 T cell function by examining production of granzyme B on CD8 T cells (Figure 5D). Indeed, the frequency of granzyme B-producing CD8+ T cells was the highest in the triple combination therapy group (Figure 5D). Together, our results strengthened the concept that Th1 skewing response triggered by soquelitinib might lead to improved CD8 function.

### Soquelitinib treatment decreases expression of T cell exhaustion markers

Previous reports have shown that T cell exhaustion limits durability of antitumor response triggered by ICB^17^. We compared the transcription profile of TILs purified from CT26 tumors from each treatment group and found that several genes associated with T cell exhaustion were upregulated following anti-PD1 treatment (Figure 5E). This finding was in line with a previous study showing increased infiltration of exhausted T cell clones following ICB therapy^18^. Interestingly, combining anti-PD1 with soquelitinib resulted in a striking reduction in the transcript levels of several exhaustion markers, including LAG3, TIGIT, Tim3, PD1, Eomes and Helios (Figure 5E). In addition to reduced exhaustion marker expression, we also found an increase in a subset of CD8+ memory (CD44+) T cells which express high levels of PD1 and TCF-1 from tumors treated with soquelitinib (Supplementary Figure 2). Previous studies have shown that TCF-1 expression on CD8 T cells defines a subset of exhausted cells that retain proliferation and stem-like renewing potential. Collectively, these findings suggested that soquelitinib treatment reduced exhaustion and improved T cell stemness in the tumor microenvironment.

### Soquelitinib treatment reinvigorates T-cell function in vitro

To better characterize the effect of soquelitinib on T cell exhaustion, we adopted an*in vitro* culture system where T cell exhaustion gradually progresses after repeated TCR stimulation^19^. Consistent with a published report^19^, we demonstrated that OT-1 CD8+ T cells receiving multiple rounds of OVA peptide stimulation exhibited signs of T cell exhaustion as evidenced by upregulation of LAG3 and TIGIT markers (Figure 6A). Increasing concentrations of soquelitinib or DMSO were added to the culture 48 hr after the first OVA stimulation. Using this approach, we found soquelitinib inhibited the expression of LAG3 and TIGIT in a concentration dependent manner as LAG3+ or TIGIT+ OT-1 T cells were virtually undetectable in cells receiving 1μM soquelitinib. Therefore, we demonstrated that selective ITK inhibition prevented the progression of T cell exhaustion *in vitro*.

**Figure 6.**
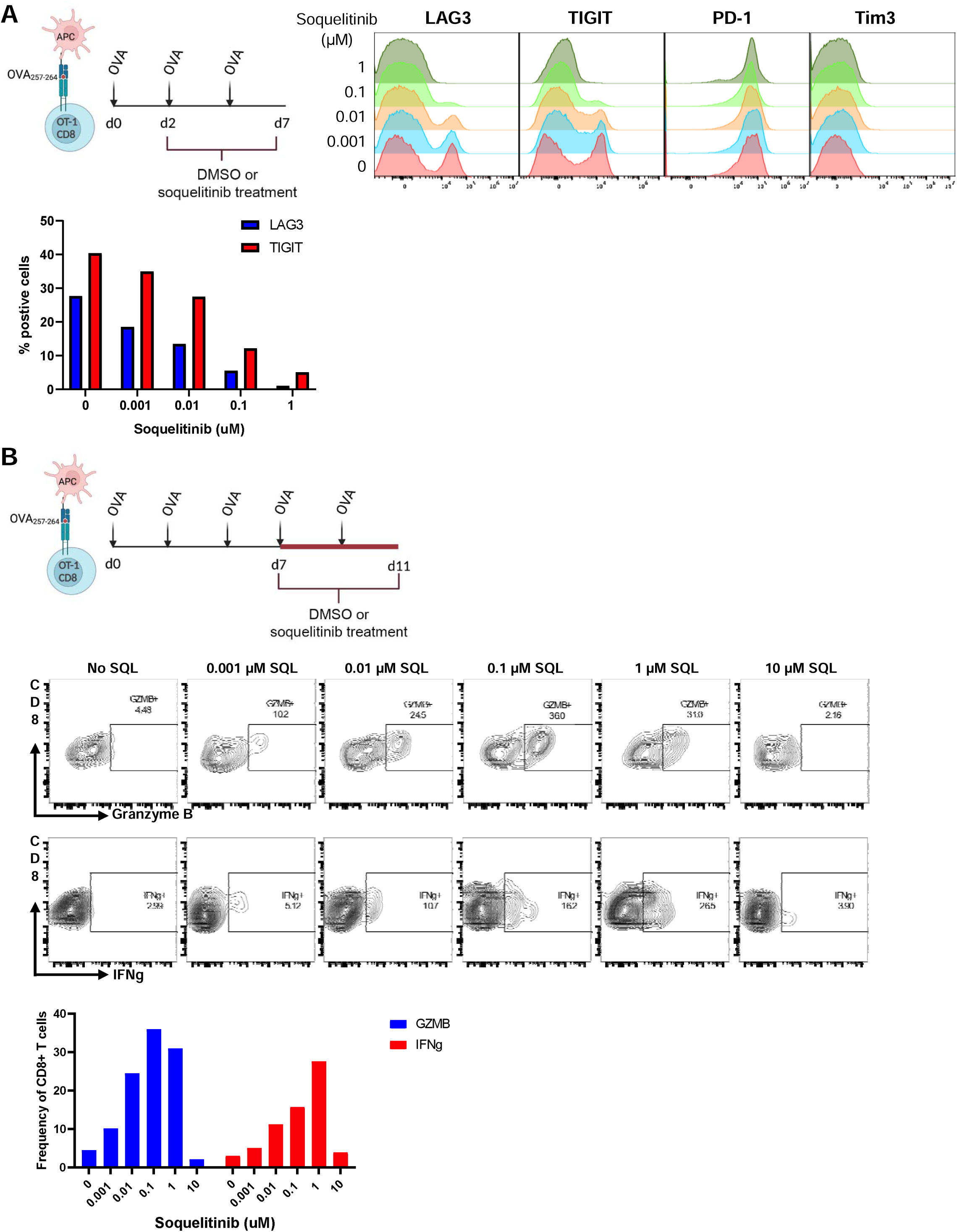
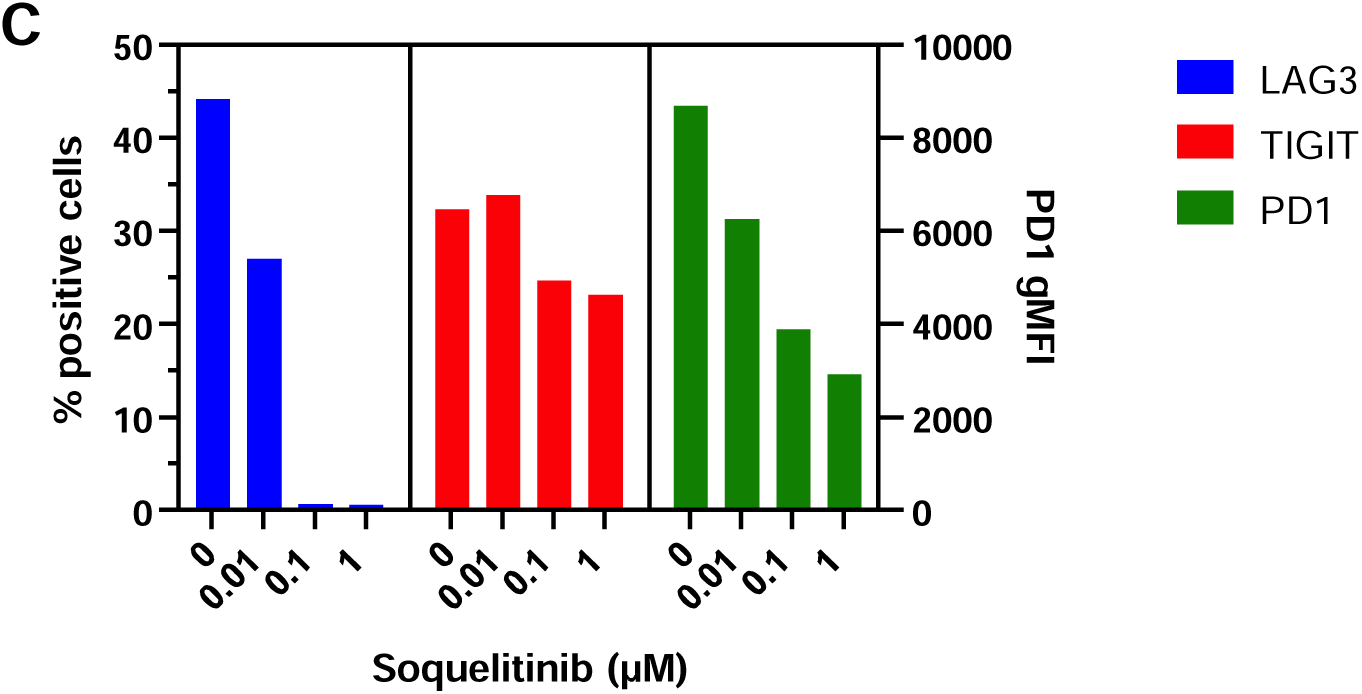
Soquelitinib inhibits T cell exhaustion process *in vitro*. (A) Schematic displaying the experimental design for evaluating the effect of soquelitinib on OT-1 T cell exhaustion process (Top left). Soquelitinib inhibits the expression of several markers associated with T cell exhaustion in a dose-dependent manner. Representative histogram overlays of indicated exhaustion marker expression in CD8+Va2+ T cells are shown for day 7 (Top right). Summary graph of two exhaustion markers, lymphocyte activation protein (LAG)3 and T cell immunoreceptor with Ig and ITIM domains (TIGIT) is shown (Bottom). (B) Schematic describing the experimental design for evaluating the effect of soquelitinib in restoring T cell exhaustion *in vitro* (Top). Soquelitinib dose-dependently (up to 1 mM) restores the potential functions of OT-1 CD8 T cells as indicated by increased production of Granzyme B and IFNg production following the 4d treatment period. Representative histograms of Granzyme B and IFNg from OT-1 T cells treated with indicated concentration of soquelitinib or DMSO control are shown (Middle). Summary graph of Granzyme B and IFNg-producing OT-1 T cells is shown (Bottom). OVA=ovalbumin; APC=antigen presenting cell; MFI=mean fluorescence intensity. SQL=soquelitinib.

We further investigated whether soquelitinib treatment might allow functional recovery of T cell exhaustion. As shown in figure 6B, we first induced T cell exhaustion in OT-1 CD8 T cells by repeated OVA stimulation for 7 days and then treated exhausted cells with either DMSO or varying concentrations of soquelitinib. As expected, we observed that T cell functions such as granzyme B and IFNγ production were greatly diminished in DMSO-treated OT-1 cells. Varying concentrations of soquelitinib restored these functions following a 4-day treatment period as indicated by increased percentage of granzyme B-and IFNγ-producing OT-1 T cells, except at the highest 10 uM concentration tested. Together, our data suggested that soquelitinib restored cytolytic function to exhausted CD8 T cells. Furthermore, we also demonstrated that repeated TCR stimulation of human CD4 T cells with anti-CD3 and anti-CD28 recapitulated the*in vitro* exhaustion process which we demonstrated in mouse OT-1 CD8 T cells. Again, soquelitinib down-regulated expression of exhaustion markers such as LAG3, TIGIT and PD1 induced by chronic stimulation through TCR in human CD4 T cells following 6-day treatment with soquelitinib. (Figure 6C).

## DISCUSSION

ITK is a member of the TEC family of kinases that plays a major role in T cell activation by integrating both TCR and CD28 costimulatory signaling pathways^9^. Studies from ITK knockout mice ^20–23^ and *in vitro* studies using shRNA knock-down ^15^ demonstrate that ITK modulates the magnitude of TCR signaling strength, leading to distinct fates during T cell differentiation and to the activation of distinct subsets of effector functions. This modulatory role for ITK is partially redundant since the structurally similar TEC kinase, RLK, has overlapping functions in Th1 cells. In this report, we describe a covalent inhibitor that is selective for ITK while sparing RLK. Our studies confirm that pharmacologic blockade of ITK by soquelitinib results in the expected inhibition of downstream signaling proteins including pPLCγ1, pERK, pS6, and NFκB and reduced secretion of IL-2. Moreover, inhibition of GATA-3 expression also was observed, supporting the known intricate relationship between ITK and GATA-3 function and the crucial role of ITK and GATA-3 in Th2 function^15^. Consistent with previously reported ITK knockout studies, we demonstrate preferential inhibition of Th2-derived cytokines such as IL-4, IL-5 and IL-13 with relative sparing of the Th1-derived cytokine, IFNγ. We find that at very high concentrations (10μM), the anti-proliferative effect of soquelitinib dominates over the effects on T cell differentiation probably as a result of inhibition of T cell receptor signaling pathways. Collectively, the selectivity and differential effects of soquelitinib on Th1 versus Th2 cells support the notion that there might be a therapeutic window for potential utility as an immunotherapy for cancer by blocking Th2 function without interfering with Th1.

Indeed, we find that soquelitinib inhibited tumor growth in several murine models including tumors that do not express ITK. In a monotherapy setting, we show increased infiltration of CD8+ T cells in both EL4 and CT26 tumor models. Furthermore, tumor-infiltrating T cells possess increased cytolytic capacity as evidenced by elevated production of IFNγ, TNF, and perforin (Figure 4). Combining soquelitinib with immune checkpoint blockade further enhances anti-tumor efficacy, leading to deep tumor regression after treatment was terminated. Several possibilities can explain this enhanced immune response to cancer in the presence of the ITK inhibitor. First, soquelitinib preferentially inhibits Th2 cytokine synthesis. Th2 cytokines have been implicated as negative regulators of the immune response to cancer^24^. There is evidence that targeting the Th2 pathway might alleviate immunosuppression in the tumor microenvironment (TME) by inhibiting the differentiation of M2-like tumor infiltrating macrophages and Th2-promoting DC subset^25^. In addition to relieving immunosuppression, selective blockade of ITK appears to modulate intratumoral balance between Th1 and Th2 cells within the TME. We show that CD4+ tumor infiltrating lymphocytes (TILs) from soquelitinib - treated tumors produce a higher percentage of cells expressing IFNγ and TNF, dominant cytokines produced by Th1 cells (Figure 4E). A previous report has shown that the Th1 cytokine, IFNγ, can induce a positive feedback loop for its own production by Th1 CD4 T cells^26^. This could potentially reinforce a bias away from a Th2-to Th1-favoring TME, resulting in further inhibition of tumor growth.

Another piece of evidence to support Th1-favoring TME was that surface expression of CXCR3, a Th1-chemokine receptor, was significantly upregulated in both CD4+ and CD8+ T cells from soquelitinib treated animals (Figure 5B,C). Our result is consistent with the study from CXCR3-deficient mice showing that CXCR3-mediated T cell recruitment to the TME is strongly associated with the induction of Th1 and cytolytic CD8 T cells^27^. Since CXCR3 upregulation is linked to a higher migratory capacity of T cells, this could potentially explain increased infiltration of CD8 T cells found in soquelitinib treated tumors.

Second, selectivity of soquelitinib to inhibit ITK over RLK enables specific blockade of ITK-mediated immune modulation, while preserving CD8 and Th1 cell functions. In support of this, additional data using a non-selective analog of soquelitinib that reacts with both ITK and RLK, indicate that treatment with an inhibitor that blocks RLK, results in marked reduction in IFNγ, IL-2 and granzyme B production in CD8 T cells (data not shown). This result mirrors the findings from the ITK-and RLK -double knockout mice in which CD8 and Th1 T cell functions were greatly impaired^28, 29^. Together, achieving significant selectivity over RLK could avoid potential impacts on vital T cell functions which are critical in driving anti-tumor immunity.

Third, preventing or reverting T cell exhaustion after soquelitinib treatment may enhance anti-tumor immunity and increase the duration of an antitumor response. It is known that continuous antigen exposure can lead to T cell exhaustion, which is characterized by gradual deterioration of CD8 T cell function and elevated expression of inhibitory receptors such as PD1, TIGIT, LAG3 and Tim3 ^30^. CD8 exhaustion has become a major barrier to cancer immunotherapy as the dysfunctional state has inverse correlation with clinical prognosis. Previous study from Weber and colleagues ^31^ has shown that transient cessation of CAR-T signaling using aSrc-family kinase inhibitor, dasatinib, that blocks proximal TCR signaling, can redirect exhausted CAR-T cells to a memory T cell state. They provided further evidence that epigenetic remodeling is required for functional reinvigoration. Since ITK is downstream of the TCR signaling pathway, it is possible that our findings are similar to findings seen with dasatinib. ITK blockade might trigger a transient rest to disrupt continuous engagement between TCR and tumor antigen without causing permanent inhibition of TCR signaling. Consistent with this notion, we find a dose-dependent reversal of the exhaustion process as evidenced by increased production of granzyme B and IFNγ in the presence of up to 1 μM concentrations of soquelitinib (Figure 6B). Future studies are needed to validate whether soquelitinib inhibits acquisition of the epigenetic hallmarks of T cell exhaustion and induces transcriptomic and epigenetic reprograming. Together, our findings show novel utility of soquelitinib in mitigating T cell exhaustion, thereby prolonging anti-tumor response.

These studies have important implications for tumor immunotherapy. Checkpoint blockade has emerged as a major therapeutic advance for cancer^32^. However, several resistance mechanisms have been elucidated including insufficient tumor immunogenicity, an immunosuppressive TME, defects in IFNγ signaling and T cell exhaustion^33^. It is possible that ITK blockade can overcome many of these resistance mechanisms and enhance immune checkpoint therapy. In fact, our murine studies demonstrate enhacement of anti-tumor activity when soquelitinib is added to anti-PD1 and anti-CTLA4 (Figure 5). Further work by Strazza et al have shown that ITK signaling is involved in the PD1-PDL1 signaling pathway, which results in inhibition of anti-tumor T cell immunity^3^. ITK blockade would be expected to prevent this inhibition and could be synergistic with anti-PD1 directed therapies.

The impact of ITK blockade on Th2 function and cytokine production raises the possibility of using an ITK inhibitor for treatment of atopic diseases. Several therapeutic agents which block Th2 cytokines or their receptors are now used in the treatment of immune diseases^34, 35^. It is possible that an ITK inhibitor would have significant impact on atopic disorders by inhibiting production of many Th2 produced cytokine*s*.

In conclusion, we have described inhibition of ITK as a novel approach to enhance the anti-tumor immune response, which occurs through several mechanisms including increased infiltration and function of cytotoxic lymphocytes and the reduction and reversal of T cell exhaustion. These observations support the use of a selective ITK inhibitor as a potential therapy for cancer. We have shown in mouse models that this approach has additive benefit to the use of recognized checkpoint inhibitors. Our*in vitro* data using human T cells are consistent with the murine studies. Furthermore, our preliminary work based on tumor biopsy in a Phase I clinical trial for T cell lymphoma is also consistent with the mouse studies (Ding, N., et.al., abstract, Hematological Oncology, in press). A potential extrapolation of our observations is that immunotherapy of cancer can be improved, while autoimmune toxicities are diminished.

## Supporting information

Supplemental table plus figures

