## Supplemental table plus figures for "Selective Inhibition of Interleukin-2 Inducible T Cell Kinase (ITK) Enhances Anti-Tumor Immunity in Association with Th1-skewing, Cytotoxic T cell Activation, and Reduced T Cell Exhaustion"

**Hsu, L-Y et.al.**

**Supplementary Material**

**Supplementary Table 1**. Unique nested chymotryptic peptide sequences present in soquelitinib treated ITK identified by deconvolution of mass spectra demonstrating an increased mass equal to the mass of soquelitinib.

| Amino Acid Sequence of Chymotrypic Fragments Bound to Soquelitinib | Corresponding Amino Acid Position in the Sequence of ITK |
| --- | --- |
| EFMEHGC[+514.170]L | 436 to 443 |
| EFMEHGC[+514.170]LSDY | 436 to 446 |
| EFMEHGC[+514.170]LSDYL | 436 to 447 |

**Supplementary Figure 1**

**A**

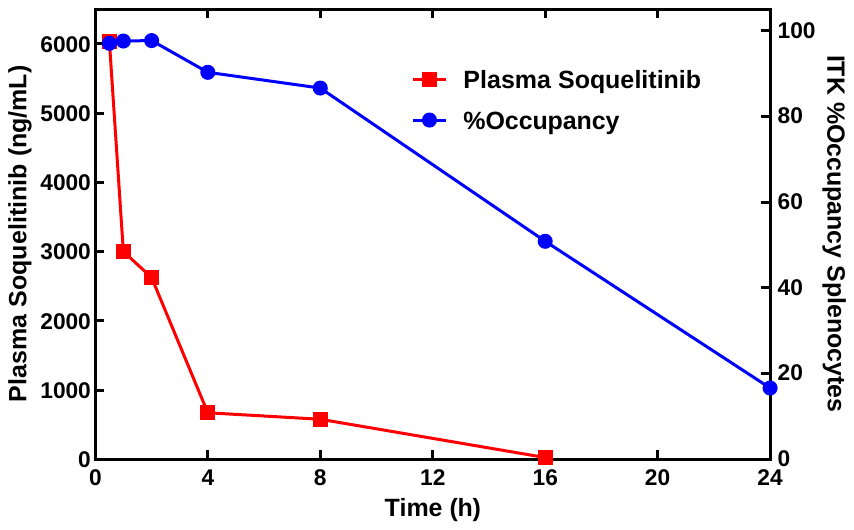

**Supplementary Figure 1. Occupancy of ITK in mouse splenocytes after oral administration.** Splenocytes and plasma were harvested at various time points from mice after administration of a single, oral dose of 50 mg/kg soquelitinib.  A competition assay was used to measure the in vivo administered drug bound to ITK.  ITK occupancy was measured in cell lysates using a biotinylated analog of soquelitinib and a fluorescent avidin detector.  Plasma levels of soquelitinib were measured using HPLC-MS/MS.

**Supplementary Figure 2**

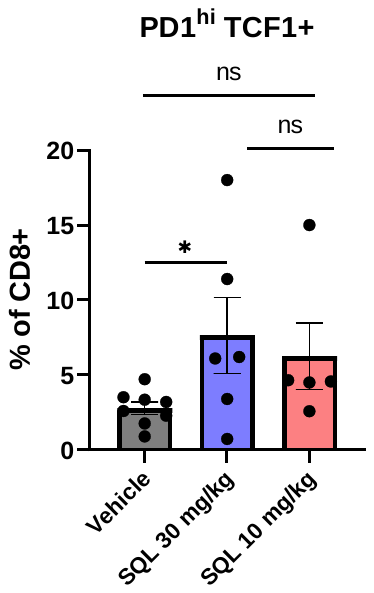

**SQL, 10 mg/kg**

**SQL, 30 mg/kg**

**Vehicle**

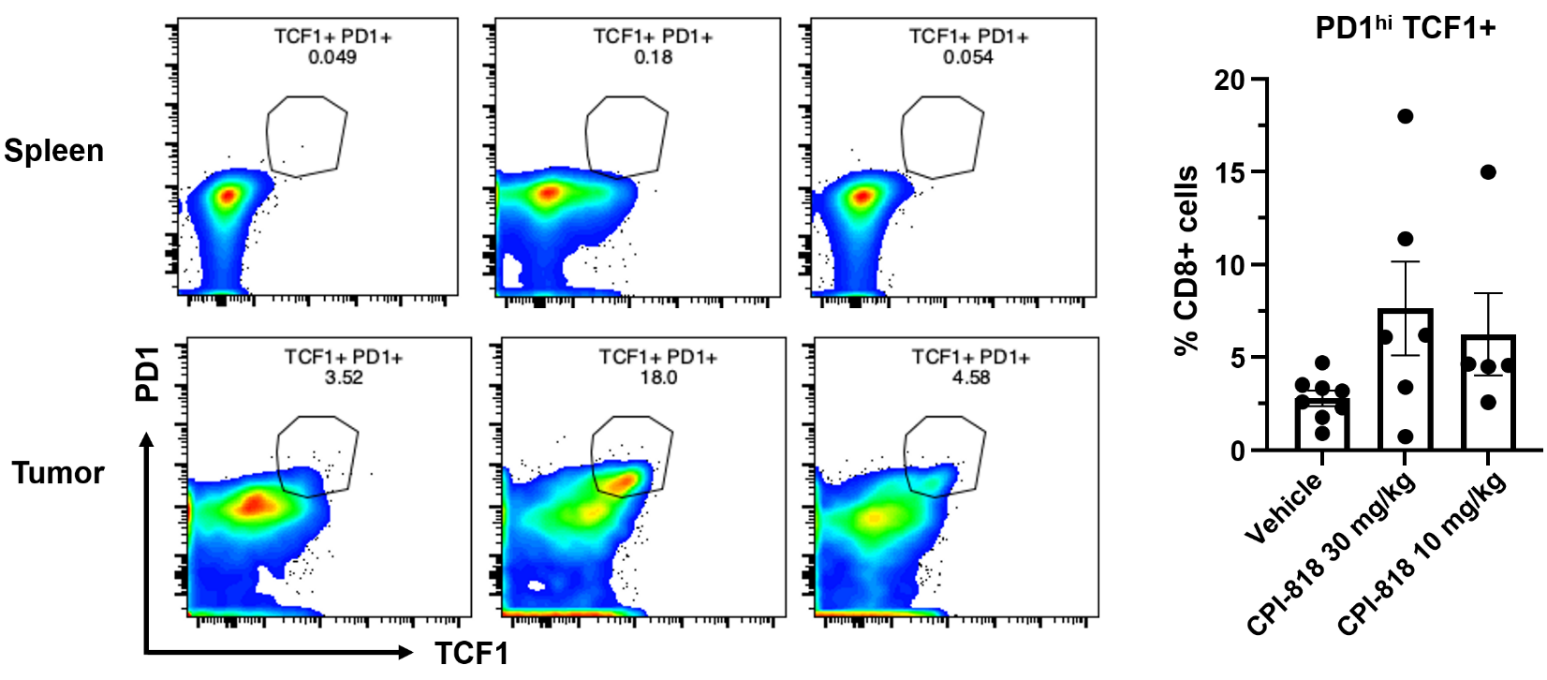

**Supplementary Figure 2. Soquelitinib increases the frequency of PD1^high^TCF1+ CD8 TILs.**

(A) Representative flow data showing the percentage of PD1^high^ T cell factor 1 (TCF1)+ CD8 T cells is increased in soquelitinib treated tumors but not in spleens (Left). Summary graph quantifying the increase in of PD1^high^TCF1+ CD8 population is shown (Right). EL4 tumor-bearing mice were treated with either vehicle or solution-formulated soquelitinib (30 mg/kg, or 10 mg/kg) for 8 days. PD1^high^TCF1+ CD8 TILs were assessed by flow cytometry. SQL= soquelitinib. NS, not significant, * p ≤ 0.05. Data displayed as means ± SEM.
